# *Deepwater Horizon* crude oil reduces aerobic capacity of birds

**DOI:** 10.1101/2022.02.02.478827

**Authors:** Christopher G. Goodchild, Jeffrey B. Krall, Arvind Santhanakrishnan, Sarah E. DuRant

**Affiliations:** Oklahoma State University, Department of Biology, Stillwater, OK 74078, USA; University of Central Oklahoma, Department of Biology, Edmond, OK 73034, USA; University of Colorado Anschutz Medical Campus, Department of Biochemistry and Molecular Genetics, Aurora, Colorado, 80045, USA; University of Arkansas, Department of Biological Sciences, Fayetteville, AR 72701, USA

**Keywords:** oil spill, avian ecology, energy budget, damage assessment

## Abstract

Crude oil spills can have catastrophic effects on marine and inland ecosystems, yet it is difficult to accurately quantify the extent of ecological damage caused by oil spills. For instance, avian population damage assessments conducted after large oil spills (e.g., *Deepwater Horizon* spill) often focus on the number of visibly oiled birds. However, birds without visible oiling can exhibit hematological damage from oil ingestion. If such hematological responses limit oxygen deliver to tissues and impair aerobic performance, then energy-mediated effects from oil ingestion may ultimately affect endpoints of demographic significance (e.g., survival and reproduction). We investigated whether oil ingestion affects aerobic performance in birds by orally dosing zebra finches (*Taeniopygia guttata*) with 2 or 6 mL/kg of weathered MC252 crude oil for 28 days. After 14 and 28 days of dosing, we measured hematological indices (oxidative damage, packed cell volume [PCV], hemoglobin, reticulocytes), maximum metabolic rate (MMR), resting metabolic rate (RMR), and short-distance flight performance. Finches exposed to oil exhibited lower hemoglobin and PCV, higher reticulocyte counts, and greater oxidative damage. Shifts in these hematological indices appeared to alter organismal energetics, resulting in reduced MMR, RMR, and aerobic scope. Short-distance burst-flight was not negatively impacted by oil ingestion. Collectively, these results suggest oil ingestion impairs metabolic performance, which may negatively impact a bird’s ability to perform sustained energetically expensive activities (e.g., migration).

**Significance:** The 2010 *Deepwater Horizon* oil spill released an unprecedented volume of crude oil (MC252) into the northern Gulf of Mexico and contaminated 2100 km of shoreline habitat that serves as critical breeding grounds and migratory stop-over sites for birds. Here we describe the impact of oil ingestion on the aerobic capacity of zebra finches, a model songbird. Oil ingestion reduced maximum metabolic rate and aerobic scope, which may be caused in part by hematological damage. These data suggest oil ingestion limits the ability of birds to perform essential energetically demanding activities (e.g., migration, nest incubation), thus quantification of avian injury based on external oiling alone may underestimate the true impact of oil spills on avian populations.

## 1. INTRODUCTION

Large marine oil spills as well as smaller but more frequent inland oil spills cause extensive ecological damage ^1–3^. For instance, the *Deepwater Horizon* (DWH) oil spill lasted 87 days, released 779 million L of Light Louisiana Crude Oil into the Gulf of Mexico in 2010, impacted 2100 km of coastline, and caused widespread oiling of shorebirds as well as songbirds inhabiting nearshore habitats ^4–6^. External exposure to crude oil damages the microstructure of feathers, which can lead to hypothermia, impaired flying performance, emaciation, and death ^7,8^. Because external oiling has direct implications for survival, avian damage assessments following large oil spills are often estimated from beach surveys of visibly oiled birds and oiled carcasses ^9–12^. According to such beach surveys, the DWH oil spill caused an estimated 102,000 bird mortalities ^13^, though some estimates suggest avian mortality was much greater ^10,14,15^. While surveys of visibly oiled birds may be predictive of mortality from external oiling, birds without visible oiling can still be exposed to crude oil internally by ingesting contaminated prey or preening oiled feathers ^16,17^. Therefore, avian damage assessments may underestimate the impact of oil spills on avian populations because beach surveys typically do not include potential toxic effects from oil ingestion.

There is robust evidence that birds orally exposed to crude oil exhibit hematological effects caused by oxidative damage to red blood cells (RBCs) from exposure to polycyclic aromatic hydrocarbons (PAHs) and PAH metabolites ^18,19^. Specifically, increased intracellular inclusion bodies (i.e., Heinz bodies), decreased packed cell volume (PCV), and higher reticulocyte (i.e., immature RBCs) counts have been observed in free-living birds inhabiting areas impacted by oil spills ^20–22^. While such shifts in hematological indices are often observed in birds that experience heavy external oiling, similar hematological responses have also been observed in birds without visible external oiling. For instance, shorebirds inhabiting areas potentially impacted by the DWH oil spill exhibited RBCs containing Heinz bodies, reduced PCV, and elevated reticulocytes, even though these birds had no visible signs of oiling ^17^. Although shifts in hematological indices have been observed in birds without external oiling, hematological effects from crude oil ingestion have yet to be linked to adverse outcomes related to avian fitness or survival.

Shifts in hematological indices caused by crude oil ingestion may contribute to a reduction in a bird’s aerobic scope, which broadly describes the rate of energy mobilization for activity and is defined as the difference between maximum metabolic rate (MMR) and basal metabolic rate (BMR)^23,24^. In endotherms, BMR is the post-absorptive resting metabolic rate of an individual within their thermoneutral zone and represents the minimum energy needed to sustain an animal. If crude oil ingestion decreases PCV, then birds may have impaired oxygen carrying capacity of blood, resulting in a reduction in MMR. Additionally, birds that have ingested crude oil may exhibit increased resting metabolic rate (RMR)—a similar metric as BMR, but assumes the resting rate measured may not be the absolute minimum metabolic rate—in response to greater detoxification and repair costs from oxidative damage to tissues ^25^. Alternatively, exposure to contaminants may cause a reduction in RMR by suppressing nonessential physiological processes in order to maintain aerobic scope for essential behaviors or to increase the amount of energy available for detoxification ^26^. Several studies have documented a reduction in PCV and increased oxidative stress in birds exposed to crude oil ^20,22,27^, yet it remains unclear whether hematological effects from crude oil ingestion translates to a reduction in MMR and/or altered RMR. Shifts in MMR and RMR in birds exposed to crude oil is ecologically relevant because a reduction in aerobic scope would have direct implications for a bird’s ability to perform energetically expensive physiological and behavioral processes important for survival, such as flying performance.

In this study, we orally dosed zebra finches (*Taeniopygia guttata*) with weathered MC252 crude oil daily for 28 days. We predicted zebra finches exposed to crude oil would exhibit RBC damage, decreased PCV and hemoglobin (Hb), and increased reticulocytes. In addition to shifts in hematological indices, we predicted zebra finches orally dosed with crude oil would exhibit a reduction in MMR and an increase in RMR. Finally, if birds orally dosed with crude oil exhibit a reduced aerobic scope, then we predicted oil-exposed birds would have impaired flying performance (see Table S1 for a summary of study predictions). The zebra finch is a commonly used model for avian toxicity testing and is particularly useful for understanding the effects of oil ingestion in songbirds inhabiting areas impacted by the DWH oil spill (e.g., seaside sparrow [*Ammodramus maritimus*]). Additionally, the effects of oil ingestion on zebra finch physiology is broadly applicable to understanding ecologically relevant energy-mediated effects in waterfowl and shorebirds exposed to crude oil.

## 2. METHODS

### 2.1. Animal husbandry and experimental design

Male zebra finches were housed in cages (30×40×40 cm) in same-sex groups of 2-3 birds to avoid distress from social isolation, and all birds within a cage received the same treatment. Female zebra finches were unavailable for this study due to logistical constraints. During the experiment, birds were fed white millet *ad lib* and had access to fresh water. The animal room was maintained on 14:10 light:dark photoperiod at 21.5±1.5 °C and 35±10% humidity. Mass, furcular fat score (range: 0-3), RBC indices, metabolic rates, and flying performance data were measured prior to exposure to crude oil and after 14 and 28 days of daily crude oil dosing. For each timepoint, we first measured flying performance, then measured RMR overnight. Birds were returned to their cages the following morning just prior to the lights turning on and given several hours to eat and drink, then we measured MMR and collected blood to measure hematological indices. In this way, blood sampling did not affect flight trials or metabolic measurements (see Fig. 1 for an overview of data collection sequence). We exposed a total of 39 zebra finches to peanut oil (control) or 2 or 6 mL/kg weathered crude oil (N=13 per group), though sample sizes varied across endpoints and over time due to limited sample volume or birds failing to complete MMR or flight trials (final sample sizes are represented in figures).

**Fig. 1:**
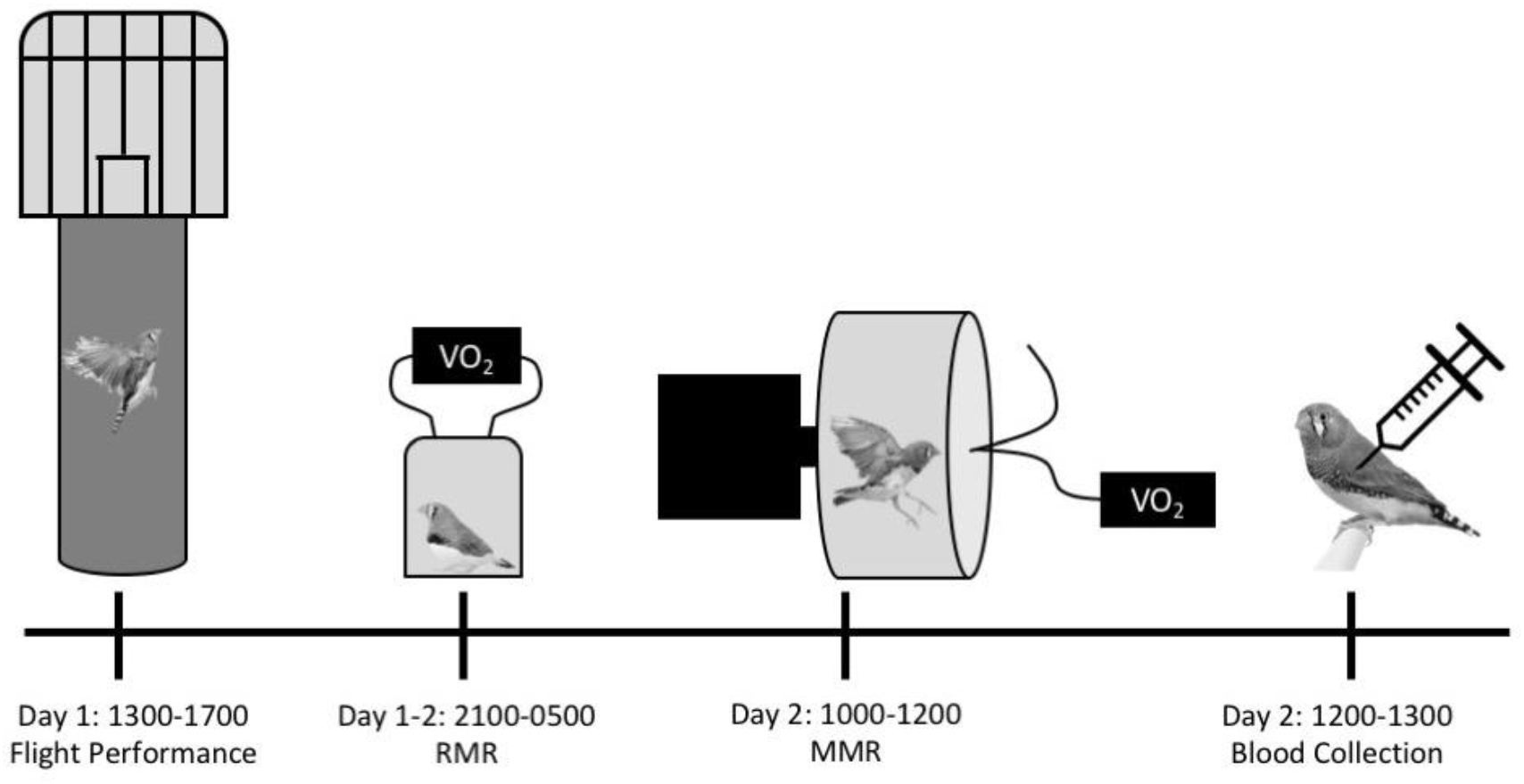
Sequence of data collection over two days that occurred prior to dosing and after14 and 28 days of dosing with peanut oil (controls) or 2 or 6 mL/kg of weathered crude oil. (RMR=resting metabolic rate measured overnight, MMR= maximum metabolic rate measured in a hop-flutter wheel.)

### 2.2. Dosing

Male finches were orally dosed daily with 2 or 6 mL/kg bird mass of artificially weathered (85% original volume; Suppl. Material^28^) MC252 crude oil (Gulf Coast Restoration Organization, Batch #: SO-20111116-MPDF-003) using a stainless-steel feeding tube. For the 2 mL/kg dose, crude oil was diluted in peanut oil, so the final delivery volume was equivalent across treatments. To reduce regurgitation, we prepared a fresh delivery matrix of peanut oil and hard-boiled egg yolk (2:3) to which we added crude oil (Fig. S1, Suppl. Material). The final delivery volume, including the egg matrix, was 20 mL/kg for all treatments and the control.

### 2.3. Hematological indices and heme degradation product formation

We collected blood (~100 μl) via venipunction of the alar vein to measure various hematological indices. We measured total Hb concentration in whole blood (~5 μl) in duplicate using the HemoCue system (Ängelholm, Sweden). To measure reticulocytes, we incubated 10 μl of whole blood in the suprarvital stain new methylene blue (Rica Chemical Company; Arlington, TX, USA) for 25 minutes prior to preparing blood smears. For each bird, a single observer (RS) who was blind to treatment examined 1000 RBCs and counted the number of reticulocytes, which were defined as RBCs with a complete reticular ring or at least five reticular clusters ^29^. We determined PCV by centrifuging whole blood in microhematocrit tubes, then measured the percent RBC *v*/*v* whole blood. We assessed oxidative damage to RBCs by measuring the formation of heme degradation products (HDPs), which are endogenous autofluorescent byproducts of oxidative damage to Hb molecules ^30,31^. HDPs were recently shown to be an indicator of oxidative damage in avian RBCs^32^. We measured HDPs according to previously established methods^32^ (Fig. S2; Suppl. Material).

### 2.4. Metabolic Performance

#### 2.4.1. Resting metabolic rate

To measure RMR, we removed birds from their cages 30 minutes after the aviary lights turned off at 2100 h and placed birds in 1 L respirometry jars supplied with a continual supply of ambient air at a flow rate of 400 mL/minute. To determine RMR, we calculated VO_2_ and VCO_2_ using equations described by Bartholomew et al. (1981), with correction for baseline drift (Suppl. Material).

#### 2.4.2. Maximal Metabolic Rate

To measure MMR, we constructed an air-tight hop-flutter wheel (diam.= 35 cm; height=14 cm) using a polypropylene cylinder with plexiglass panels affixed to both sides (*sensu* Chappell et al. 2006). Air was scrubbed of CO_2_ and water vapor using Drierite and pumped into the hop-flutter wheel at a flow rate of 4200 mL/minute. We monitored %O_2_ in excurrent air in real-time for exercised birds until the O_2_ consumption profile plateaued and further increasing the rotational speed of the hop-flutter chamber did not increase O_2_ consumption, at which point we turned off the hop-flutter wheel (Suppl. Material). All birds were visibly panting after the hop-flutter wheel was turned off. We calculated aerobic scope as the difference between MMR and RMR.

### 2.5. Flying performance

To measure short-distance flight speed, we constructed a vertical flight chamber similar to Kullberg et al. (2020)^35^ (Suppl. Material). Prior to the experiment, birds were trained to fly in the vertical flight chamber and only birds that consistently completed their first attempt on the final day of training were included in the experiment. There were no differences in tarsus length, body width, or wing chord or length among finches (*F*<2.25, *p*>0.14; Table S2). During the experiment, each bird was given 7 attempts to complete 4 flights for each day of data collection. Birds that failed to complete four flights were returned to their cage (see section 2.7 for how success/fail attempts were handled). Throughout training and the experiment, a single researcher (JBK) placed birds on the perch at the bottom of the flight chamber in an effort to maintain consistent handling effects.

Each vertical flight attempt was video-recorded using a Canon camcorder (Vixia HF R700) that captured video at 30 frames per second (fps) and was positioned 4 m from the flight chamber. Videos of each bird’s vertical flight were analyzed using ImageJ, whereby we converted videos to individual frames and digitally measured flight distance. Vertical speed was calculated by dividing the measured linear distance traveled by the time it to took a bird to travel the respective distance (Suppl. Material).

### 2.6. PAH quantification

Analysis of Σ50 PAHs in artificially weathered crude oil was performed by ALS Environmental (Kelso, WA, USA). The analysis of PAHs, alkyl PAH homologues, and related hetero-compounds was conducted using gas chromatography (GC) with low-resolution mass spectrometry (MS) using selected ion monitoring based on USEPA method 8270D. The minimum detection limit (MDL) was 4 mg/kg. Σ50 PAHs in weathered crude oil used in this study was 12,942 mg/kg (Fig. 2; Table S3), which is comparable to previous studies (13,808 mg/kg)^28^.

**Fig. 2:**
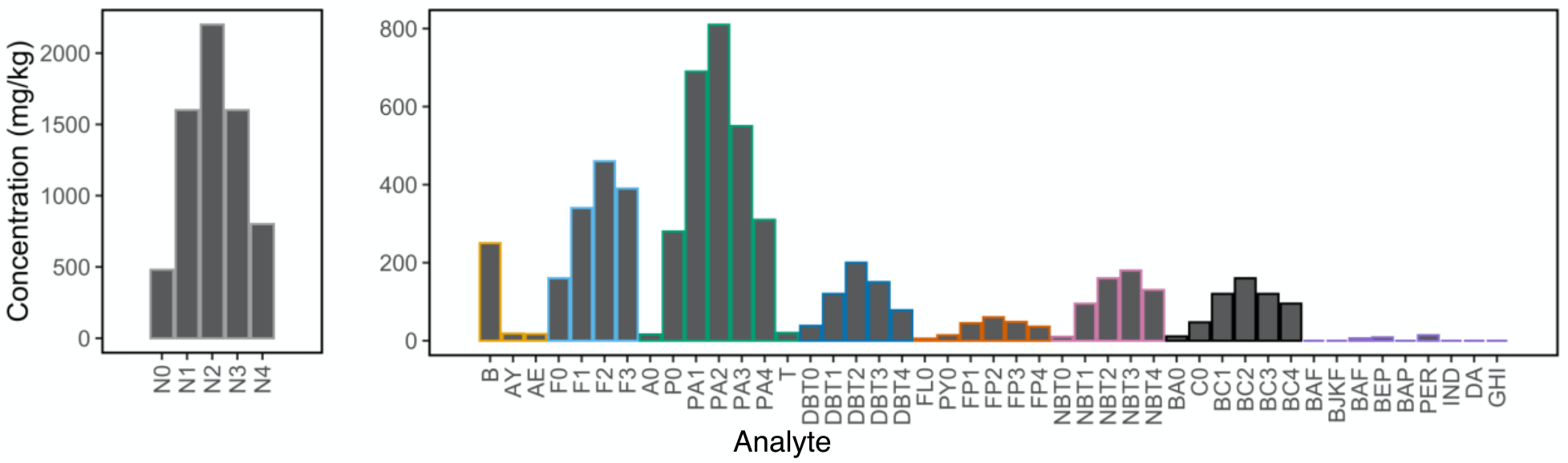
Polycyclic aromatic hydrocarbons present in weathered crude oil. See Supplemental Information, Table S3 for definitions of analyte abbreviations.

### 2.7. Statistics

Fat score, hematological, metabolic, and flight data were analyzed with linear mixed models (LMM) using the ‘nlme’ package in R ^36^(Version 3.6.3), with treatment and time as fixed effects and individual as a repeated random effect. For each model, we used the ‘car’ package to calculate analysis of deviance for each model to examine Wald **χ**^2^ of main effects and interaction terms ^37^. All models included a treatment×time interaction, except fat score which we re-ran without the interaction because it was not significant (treatment×time: **χ**^2^=3.03, *p*=0.55). Mass was included as a fixed effect covariate for LMMs of fat score, RMR, MMR, aerobic scope, and take-off velocity and acceleration. Additionally, mass was initially included in the LMM vertical flight speed but was not a significant covariate (**χ**^2^=1.84, *p*=0.18), for this reason we re-ran the LMM without mass as a covariate. We also examined the effects of crude oil dosing on flight speed by analyzing % change in flight speed on days 14 and 28 relative to each bird’s flight speed on day 0 using generalized linear models (GLMs). To test for treatment difference in the time until the first failed flight attempt, we used a cox proportional hazard model in the ‘survival’ package in R.

## 3. RESULTS

### 3.1. Mass, fat score, and morphology

Regardless of treatment, mass decreased over the course of the experiment (LMM: time: **χ**^2^=24.92, *p*<0.001; treatment×time: **χ**^2^=4.43, *p*=0.35; treatment: **χ**^2^=1.73, *p*=0.42). Specifically, mass was lower on day 14 (14.82±1.31 g [mean±std. dev.]; |t|=4.31, *p*<0.001) and day 28 (15.14±1.50 g [mean±std. dev.]; |t|=2.47, *p*=0.02) compared to day 0 (15.52±1.67 g [mean±std. dev.]). Fat score was consistent over time (time: **χ**^2^=3.45, *p*=0.18), but varied by treatment (Fig. S3; LMM: treatment: **χ**^2^= 9.58, *p*=0.01), which appeared to be driven by birds in 6 mL/kg crude oil treatment having a lower fat score compared to controls (|t|=3.06, *p*=0.004), whereas there was no difference between birds exposed to 2mL/kg crude oil and the controls (|t|=1.89, *p*=0.07). Further, heavier birds tended to have more fat than smaller birds (mass: **χ**^2^=37.50, *p*<0.001).

### 3.2. Hematological indices

Birds exposed to crude oil exhibited a lower PCV over time (Fig. 3A; LMM: treatment×time: **χ**^2^=24.66, *p*<0.001; treatment: **χ**^2^=14.25, *p*<0.001 time: **χ**^2^=2.75, *p*=0.25). Specifically, birds exposed to 2 mL/kg or 6 mL/kg crude oil exhibited a decrease in PCV on both day 14 (2 mL/kg: |t|=2.35, *p*=0.02; 6 mL/kg: |t|=3.19, *p*=0.002) and day 28 (2 mL/kg: |t|=3.46, *p*=0.001; 6 mL/kg: |t|=4.60, *p*<0.001) compared to the controls. Total Hb concentration also decreased over time in birds exposed to crude oil (Fig. 3B; LMM: treatment×time: **χ**^2^=38.56, *p*<0.001; treatment: **χ**^2^=4.80, *p*=0.09; time: **χ**^2^=7.57, *p*=0.02). Birds exposed to 6 mL/kg crude oil exhibited a decrease in Hb concentration on day 14 compared to controls (|t|=4.82, p<0.001). Similarly, birds exposed to 2 mL/kg appeared to have lower Hb concentration compared to controls on day 14 (|t|=1.75, *p*=0.08). Birds in both oil treatments exhibited lower Hb concentrations compared to the controls on day 28 (2 mL/kg: |t|=3.35, *p*=0.001; 6 mL/kg: |t|=5.54, p<0.001). Reticulocytes increased over time in birds that ingested crude oil (Fig. 3C; LMM: treatment×time: **χ**^2^=24.46, *p*<0.001; treatment: **χ**^2^=26.50, *p*<0.001; time: **χ**^2^=59.59, *p*<0.001). Birds exposed to 2 mL/kg and 6 mL/kg crude oil exhibited an increase in the number of reticulocytes on both day 14 (2 mL/kg: |t|=2.32, *p*=0.03; 6 mL/kg: |t|=2.31, *p*=0.03) and day 28 (2 mL/kg: |t|=4.64, *p*<0.001; 6 mL/kg: |t|=3.77, *p*<0.001) compared to control birds.

**Fig. 3:**
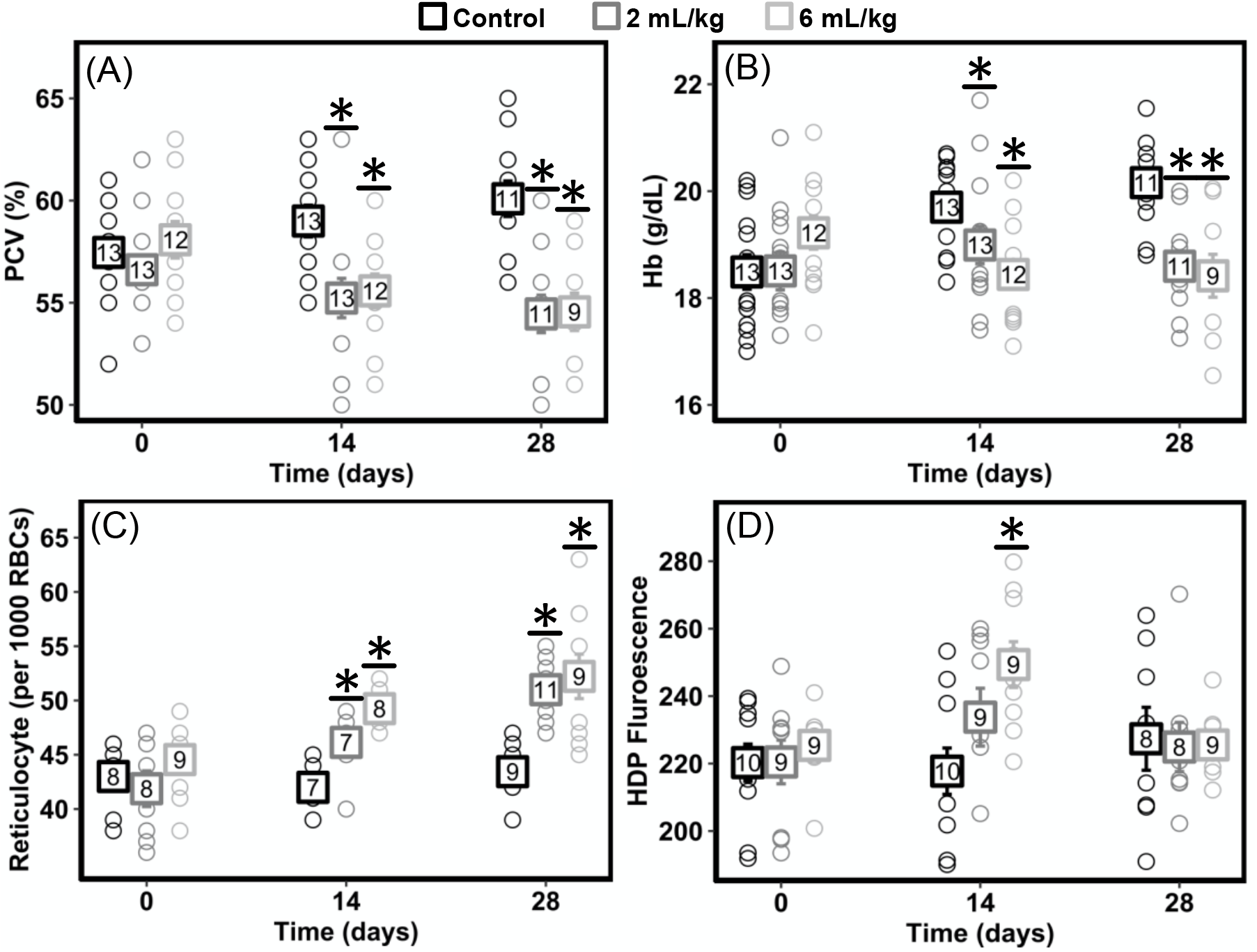
Hematological indices from zebra finches prior to dosing (day 0) and after 14 and 28 days of oral dosing with peanut oil (controls) or weathered crude oil. (A) packed cell volume (PCV); (B) total hemoglobin (Hb) concentration; (C) reticulocytes; (D) oxidative damage measured by heme degradation product (HDP) fluorescence in red blood cells (squares and whiskers represent mean±std. error, numbers within squares represent sample sizes, open circles represent values for individual birds, asterisks denote significant difference [*p*<0.05] compared to the control).

Birds exposed to crude oil appeared to vary in HDP fluorescence over time (Fig. 3D; LMM: treatment×time: **χ**^2^=8.88, *p*=0.06; treatment: **χ**^2^=3.95, *p*=0.14; time: **χ**^2^=5.44, *p*=0.07). Further, pairwise comparisons suggest that birds exposed to 6 mL/kg exhibited increased HDP fluorescence on day 14 compared to controls (|t|=2.26, *p*=0.03). HDP fluorescence in birds exposed to 6 mL/kg was not different than the controls on day 28 (|t|=0.59, *p*=0.56). There were no differences between birds exposed to 2 mL/kg crude oil and controls on days 14 or 28 (for all comparisons |t|<1.35, *p*>0.18).

### 3.3. Metabolic rate

Birds exposed to crude oil exhibited a decrease in MMR over time (Fig. 4A; LMM: treatment×time: **χ**^2^=14.93, *p*=0.005; treatment: **χ**^2^=3.14, *p*=0.21; time: **χ**^2^=6.28, *p*=0.04). Specifically, birds exposed to 6 mL/kg crude oil exhibited a decrease in MMR on days 14 and 28 compared to the controls (day 14: |t|=2.41, *p*=0.02; day 28: |t|=3.12, *p*=0.002), whereas there were no differences in MMR for birds exposed to 2 mL/kg crude oil compared to controls over time (day 14: |t|=1.81, *p*=0.08; day 28: |t|=0.45, *p*=0.65). Birds exposed to crude oil also exhibited a decrease in RMR over time (Fig. 4B; LMM: treatment×time: **χ**^2^=15.38, *p*=0.004; treatment: **χ**^2^=11.12, *p*=0.004; time: **χ**^2^=1.51, *p*=0.47). Specifically, birds exposed to 2 mL/kg or 6 mL/kg decreased RMR on day 14 (2 mL/kg: |t|=2.31, *p*=0.02; 6 mL/kg: |t|=3.72, *p*<0.001) and day 28 (2 mL/kg: |t|=2.05, *p*=0.045; 6 mL/kg: |t|=2.45, *p*=0.02) compared to controls. MMR and RMR of birds increased with increasing body mass (MMR: |t|=21.43, p<0.001; RMR: |t|=13.71, *p*<0.001)

**Fig. 4:**
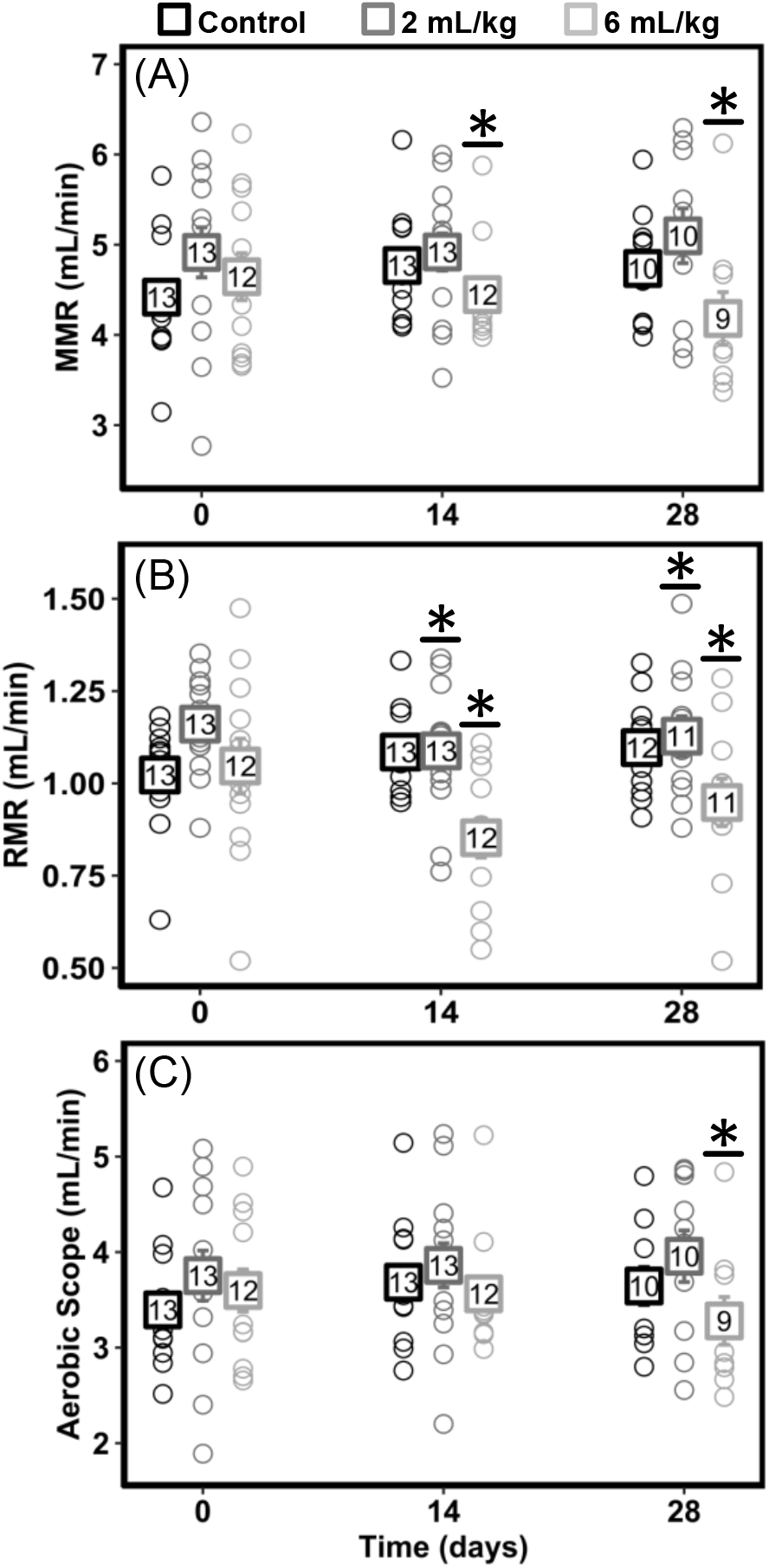
Metabolic profiles of zebra finches prior to dosing (day 0) and after 14 and 28 days of oral dosing with peanut oil (controls) or weathered crude oil: (A) maximum metabolic rate (MMR); (B) resting metabolic rate (RMR); (C) aerobic scope calculated as the difference between MMR and RMR (squares and whiskers represent mean±std. error, numbers within squares represent sample sizes, open circles represent values for individual birds, asterisks denote significant difference [*p*<0.05] compared to the control).

Birds exposed to crude oil also exhibited a significant reduction in aerobic scope over time (Fig. 4C; LMM: treatment×time: **χ**^2^=9.23, *p*=0.06; treatment: **χ**^2^=1.26, *p*=0.53; time: **χ**^2^=7.85, *p*=0.02). This trend appeared to be driven by birds in the 6 mL/kg crude oil treatment, which exhibited a decrease in aerobic scope on day 28 compared to controls (|t|=2.46, *p*=0.02), but had similar aerobic scope as controls on day 14 (|t|=1.52, *p*=0.14). There was no difference in aerobic scope in birds exposed to 2 mL/kg over time compared to controls (for both days 14 and 28: |t|<0.79, *p*>0.43). Aerobic scope of birds increased with increasing body mass (|t|=14.12, *p*<0.001).

### 3.4. Flying performance

Although not significant, vertical flight speed appeared to increase in birds exposed to crude oil over time (Fig. 5A; LMM: treatment×time: **χ**^2^=8.25, *p*=0.08; time: **χ**^2^=2.30, *p*=0.32; treatment **χ**^2^=0.91, *p*=0.64). Given the pattern in the data, we more closely examined the treatment×time trend and found that birds in the 6 mL/kg crude oil treatment appeared to have greater vertical flight speed on day 14 (|t|=1.94, *p*=0.06) and day 28 (|t|=1.98, *p*=0.05) compared to controls. Birds exposed to 6 mL/kg increased flight speed by 18% on day 14 (GLM: |t|=2.06, *p*=0.047) and exhibited a similar but non-significant 11% increase on day 28 (GLM: |t|=1.84, *p*=0.08) compared to day 0, but there were no differences in % change in flight speed for birds exposed to 2 mL/kg crude oil (Fig. 5B; for both days 14 and 28: |t|<0.32, *p*>0.75). Birds that ingested either 2 or 6 mL/kg crude oil appeared to be more likely to fail to complete four vertical flights; however, we were unable to detect a significant difference between treatments with a cox proportional hazard model (Fig. 5C; *p*=0.21).

**Fig. 5:**
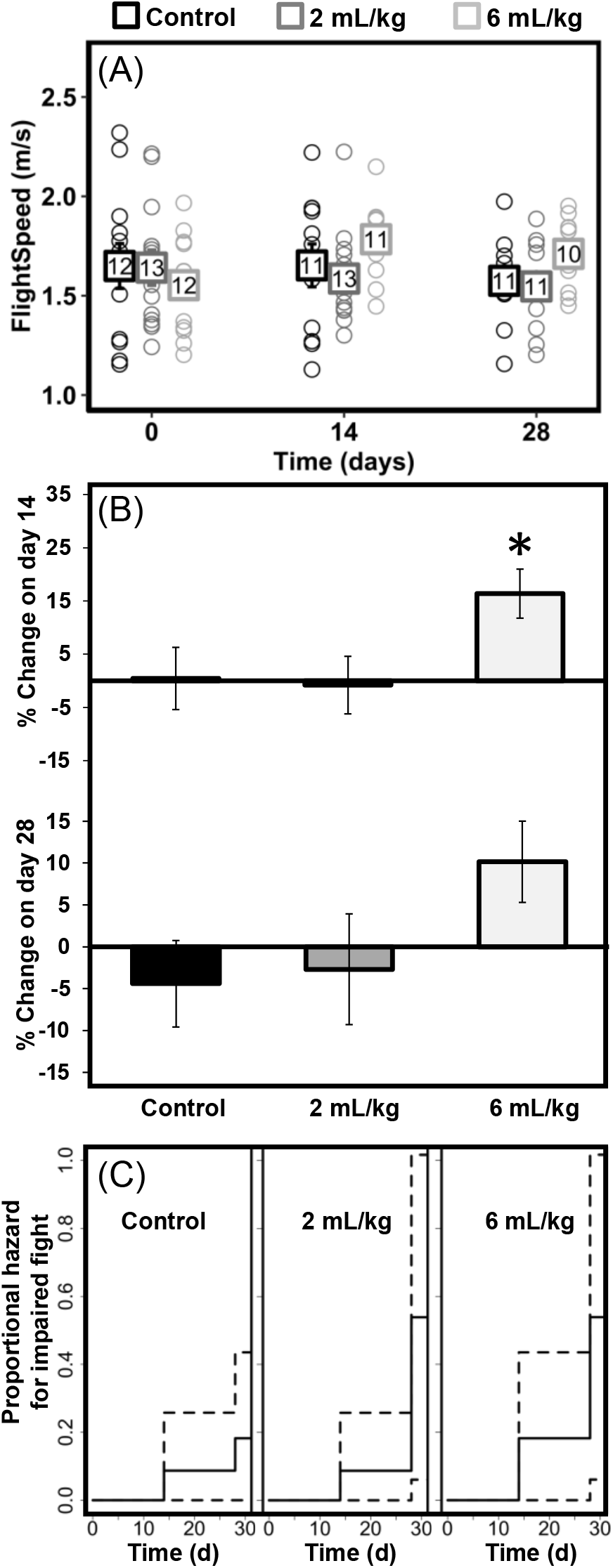
Vertical flight performance of zebra finches prior to dosing (day 0) and after 14 and 28 days of oral dosing with peanut oil (controls) or weathered crude oil: (A) Vertical flight speed (squares and whiskers represent mean±std. error, numbers within squares represent sample sizes, open circles represent values for individual birds); (B) % change relative to day 0 speed (asterisks denote significant difference [*p*<0.05] compared to the control); (C) solid line is the proportional hazard of failing the flight challenge and dashed lines are 95% confidence intervals.

## 4. DISCUSSION

Erythroid damage has been documented in diverse taxa exposed to crude oil (e.g., Leighton 1985; Alkindi et al. 1996; Da Silva Júnior et al. 2013), raising the concern for oil-exposed animals to have impaired oxygen delivery to tissue and reduced aerobic performance. Nonetheless, due to a paucity of data examining the relationships between RBC damage and whole-animal toxicity, sublethal effects from crude oil ingestion are rarely incorporated into population damage assessments for wildlife, including birds. In this study, zebra finches orally dosed with crude oil exhibited oxidative damage to RBCs and hematological responses similar to effects previously observed in free-living birds exposed to oil spills (i.e., lower PCV, higher reticulocyte counts). Additionally, crude oil ingestion reduced RMR and to a greater extent MMR, which resulted in lower aerobic scope. However, we did not detect a negative effect on flying performance (discussed below). In addition to effects on hematological indices and metabolic performance, we also found evidence of depleted fat reserves (i.e., lower fat score) in birds exposed to 6 mL/kg crude oil. While birds with a leaner body composition may exhibit increased flight speed for short distance flights ^41^, an inability to maintain fat reserves would likely limit performance during sustained long-distance flight (discussed below) and other energetically-expensive processes (e.g., reproduction). Taken as a whole, these data suggest that crude oil ingestion restructures a bird’s energy budget and may impair a bird’s ability to perform sustained energetically demanding activities.

### Effects of crude oil ingestion on hematological indices and RBC damage

This study provides further evidence of RBCs being a sensitive target for PAHs and other oxidizing chemicals present in crude oil. However, although the differences in PCV, Hb, and reticulocytes were statistically significant, the effect size was relatively small compared to findings in previous studies. For instance, the largest difference in PCV occurred on day 28, when PCV was 54% in the 6 mL/kg treatment compared to 60% in the control group. It is worth emphasizing that birds exposed to 6 mL/kg crude oil did not have pathologically low PCV, even though there were clear differences between treatments. Nonetheless, we chose moderate (6 mL/kg) and low (2 mL/kg) doses compared to other studies that orally dosed birds with 10-20 mL/kg of weathered MC252 crude oil (e.g., Cunningham et al. 2017; Dean et al. 2017; Horak et al. 2017) to better reflect concentrations birds may encounter in the wild. Furthermore, previous studies detected plasma ΣPAH concentrations >150 ng/mL in oiled seabirds ^45^. However, in a separate experiment, we dosed zebra finches with 10 mL/kg of crude oil for 14 consecutive days but were unable to detect PAHs in whole blood ^46^. Therefore, it is noteworthy that there was a shift in several hematological indices given the comparatively low doses administered in the present study. While further research is required to understand oil ingestion rates and PAH metabolism in birds exposed to oil spills *in situ* ^16^, these results suggest RBC damage is possible at lower blood-PAH concentrations.

In addition to shifts in commonly measured hematological variables, we also found that HDP fluorescence was higher in birds exposed to 6 mL/kg after 14 days of dosing but there was no difference in HDP fluorescence between the control and treatments after 28 days of dosing. Interestingly, the temporal response of HDP fluorescence differed from other hematological responses. This may be attributable to the removal of damaged RBCs from circulation, which would contribute to birds exposed to oil exhibiting a lower PCV on day 28. Furthermore, we observed higher reticulocyte counts in birds exposed to 2 or 6 mL/kg on day 28 compared to controls, suggesting damaged RBCs were being replaced with immature RBCs. Although further investigation is required to understand the biochemical and physiological relationships between HDP formation and commonly measured hematological indices, these data suggest HDP formation may be an additional tool for early detection of RBC damage before pathological conditions develop (i.e., hemolytic anemia).

An increase in HDP fluorescence in avian RBCs is consistent with previous studies that found exposure to crude oil causes oxidation of Hb, leading to the formation of Heinz bodies ^20^. The presence of Heinz bodies, caused by the aggregation of denatured Hb molecules, has been suggested as an indicator of crude oil ingestion in birds ^45,47,48^. Avian RBCs expressing Heinz bodies are typically removed from circulation or partially phagocytized in the spleen, which can lead to a pathologically low blood-oxygen level known as ‘Heinz body anemia’. However, there is considerable interspecific variation in whether and to what extent RBCs form Heinz bodies in birds that have ingested oil ^17,42^. For instance, in a previous study, we were unable to detect Heinz bodies by light or scanning electron microscopy in zebra finches orally dosed for 14 consecutive days with artificially weathered (85% original volume) MC252 crude oil^49^ (also see ^42^). An increase in HDP fluorescence in zebra finches orally dose with crude oil suggests Hb oxidation is still a concern, even if Heinz bodies are absent.

### Effects of crude oil ingestion on metabolic performance

We did not specifically measure the oxygen carrying capacity of blood, but a drop in PCV and Hb concentration suggests reduced ability of blood to transport oxygen. Consequently, depressed MMR is likely due in part to decreased PCV and Hb. However, it is unlikely lower PCV was the only factor contributing to a reduction in MMR since PCV was not pathologically low in finches exposed to crude oil. In this study, we focused on oxidative damage to RBCs, but oxidative damage likely occurred in other tissues as well ^19,50^. Thus, a reduction in MMR is likely attributable to oxidative damage to several tissues that collectively limited metabolic activity during exercise. For instance, double-crested cormorants (*Phalacrocorax auritus*) dermally exposed to crude oil had distensible hearts with flaccid musculature ^51^, which may be caused by oxidative damage to cardiomyocytes ^52^. Given that MMR is partially a function of cardiac output (Bishop 1999), oil-induced damage to heart tissue is likely an important factor contributing to a reduction in MMR. The potential for crude oil ingestion to cause cardiotoxicity is consistent with studies indicating that exposure to PAHs *in ovo* reduced avian embryonic heart rate ^53^, as well as extensive evidence of cardiotoxicity in other taxa ^54^. To understand the mechanisms contributing to reduced MMR, further research should be directed towards exploring the effects of oil ingestion on cardiac performance in vertebrates.

In addition to depressed MMR, we also observed a reduction in RMR. To meet the metabolic demands of detoxification and repair, animals can either elevate their RMR or redirect metabolic resources away from nonessential physiological processes and toward detoxification and repair^26^. In broader terms, tradeoffs between self-maintenance and other physiological processes (e.g., reproduction, immune function, molt) have been well documented in birds ^55–57^. Moreover, in a separate study, zebra finches orally dosed with artificially weathered MC252 crude oil (3.3 or 10 mL/kg) exhibited reduced activity and suppressed immune function ^46^, suggesting songbirds exposed to crude oil restructure their energy budgets. Although this is the first study to examine the effects of crude oil ingestion on RMR in songbirds, similar trends occurred in studies where birds were exposed to crude oil externally. For instance, western sandpipers (*Calidris mauri*) externally exposed to oil trended towards lower RMR (*p*=0.08)^58^. Additionally, similar effects on RMR have been observed in other taxa (e.g., fish) ^59^. It is worth noting that several studies have found that external oiling caused the opposite response to occur, whereby oiled birds increased RMR^22^ to maintain body temperature as a compensatory response to external oiling damaging feather integrity and reducing insulation ^60,61^. The differences in RMR responses highlights the potential for external and internal exposure to have substantially different effects on metabolic processes. Nonetheless, taken as a whole, these data indicate that crude oil ingestion necessitates a restructuring of a bird’s energy budget to compensate for the energetic demand of detoxification and repair.

Although there was a reduction in RMR in birds exposed to 6 mL/kg of crude oil, birds in this treatment still exhibited a reduction in aerobic scope on day 28. A reduction in aerobic scope in birds exposed to crude oil may indirectly affect ecologically important physiological processes (e.g., flying performance, immune function, parental behavior) and may ultimately affect endpoints of demographic significance, such as reduced survival and reproduction ^62–65^. Indeed, metabolic rate during flight can be up to 30× RMR ^66^. Because flight is central to migration, predator avoidance, and foraging strategies ^67^, a reduction in aerobic scope in birds that have ingested crude oil has direct implications for survival. Several previous studies have found that external exposure to crude oil impairs flying performance caused by damage to feathers ^68^, whereas results from the present study indicate flight performance may also be affected by a reduction in aerobic scope. That is, birds that ingest crude oil but do not experience external oiling are still at risk of experiencing reduced flight speed and being quicker to fatigue due to reduced MMR and aerobic scope, respectively.

### Effects of crude oil ingestion on take-off flight performance

Aerobic scope and MMR are closely linked to locomotor performance across taxa^26^, and we predicted a reduction in aerobic scope in birds exposed to crude oil would result in a concomitant reduction in short-distance flight performance. However, the opposite effect occurred: birds exposed to 6 mL/kg of crude oil exhibited increased flight speed on day 28. This response may be attributable to burst flight over a short distance being a mostly anaerobic process that does not require sustained oxygen supply to muscles for energy mobilization. Additionally, we found that birds exposed to 6 mL/kg crude oil had an overall lower fat score, which would result in a leaner body composition and may explain why birds exposed to 6 mL/kg oil exhibited increased flight speed ^41^. Although leaner body composition may facilitate increased flight speed in the short-term, a reduction in fat score is not adaptive for long-distance migratory flight. A reduction in fat score indicates depleted energy reserves and further supports that crude oil ingestion alters the energy budget of birds. Importantly, due to the sustained metabolic demand during migration, birds will engage in bouts of hyperphagia to rapidly increase mass and replenish fat reserves ^69^. Thus, if oil exposure impairs a bird’s ability to maintain fat stores, exposure to crude oil during migration may have severe consequences for a bird’s ability successfully arrive at summer breeding grounds ^70^. Indeed, other studies found externally oiled birds have impaired pre-migratory fueling and lose more weight during migration ^71–73^. Therefore, although we did not detect effects of crude oil ingestion on short-distance flights, a reduction in aerobic scope and depleted fat reserves will likely limit a bird’s ability to conduct long-distance flights ^72,74^.

## 5. CONCLUSION

We found evidence that low-level crude oil ingestion causes a restructuring of a bird’s energy budget. A particular concern highlighted by this study is the ability of crude oil to reduce aerobic scope, given that reduced aerobic scope has broad implications for numerous physiological processes. These results suggest that damage assessments that rely exclusively on surveys of visibly oiled animals may underestimate the extent to which crude oil spills impact wildlife populations because they do not consider the toxic effects of crude oil ingestion.

## ACKNOWLEDGEMENTS

The authors acknowledge Ridley Shelby, Kevin Grisham, and Weston Perrine for their assistance with data collection, and Stan Goodchild for his assistance with the construction of the vertical flight chamber. We thank Jason Belden, Jennifer Grindstaff, and Matteo Minghetti for their thoughtful critiques of the experimental design and manuscript. Christopher Goodchild received funding support from the Oklahoma State University (OSU) Interdisciplinary Toxicology Program and OSU Robberson Summer Dissertation Scholarship. Jeffrey Krall received support from the OSU Niblack Scholarship and Goldwater Scholarship.

## DATA SHARING PLANS

All data and code will be made available on a public data depository upon acceptance.

## Supplemental Methods

### Artificial weathering of crude oil

Weathering of spilled crude oil can occur by various physical and chemical processes, including biodegradation, evaporation, dissolution, emulsification/dispersion, and photo-oxidation^1^. This weathering process resulted in the chemical composition surfaces slicks in the Gulf of Mexico being chemically distinct from source oil. To better assess adverse outcomes from oil ingestion, we artificially weathered MC252 crude oil by heating 400 mL of oil to 95 °C using a digital hot plate. The oil was stirred continuously at a speed that would mix the oil without aerating it until the oil was 85% the original volume^2^. PAH concentrations in artificially weathered crude oil were quantified at ALS Environmental (Kelso, WA, USA; Fig. 2; Table S3).

### Dosing

Regurgitation of crude oil is often observed in avian toxicity tests. To reduce regurgitation, we prepared a fresh delivery matrix of peanut oil and hard-boiled egg yolk (2:3) daily to which we added crude oil or peanut oil (control). Hard-boiled egg yolk is a nutritional supplement for zebra finches. We assessed whether the delivery of crude oil within an egg-matrix reduced regurgitation by dosing birds with 3 mL/kg of crude oil either straight or within an egg matrix (N=5 per treatment), and monitored birds individually for 60 min. The egg matrix significantly reduced time till regurgitation (Fig. S1; t-test: t= 4.83, p-value = 0.001). All birds exposed to straight oil regurgitated in < 35 min, whereas only 2 birds regurgitated in less than 60 min in birds dosed with oil in an egg matrix. Furthermore, the amount of regurgitated material was lower in birds dosed with oil in the egg matrix (personal observation).

### Heme degradation product fluorescence

We measured the fluorescence of heme degradation products in whole blood as described by Goodchild and DuRant (2020)^3^. Because hemoglobin can quench the fluorescent signal produced by heme degradation products^4^, we measured hemoglobin concentration, as described elsewhere, and diluted all blood samples with deionized water to a final hemoglobin concentration of 1 mg/dL^5^. Diluted RBCs were vortexed for 10 s to lyse cells and stored at −80 C until analysis. The fluorescent intensity of HDPs does not differ between fresh blood and frozen samples (unpublished data). Immediately prior to analyzing HDP fluorescence, diluted blood samples were thawed, vortexed, and 150 μl aliquots were transferred to a flat-bottom 96-well plate (5 technical replicates per biological sample; Greiner Bio-One). We measured the fluorescence of heme degradation products in five technical replicates on a plate reader (Cytation 5, BioTek) at an excitation/emission wavelengths of 321/450 nm ^3,6^. Interassay CV was 3.71% and mean intraassay CV was 4.29% (range: 1.75-8.50%). Representative emission spectra is provided for a single individual prior to exposure (day 0) and after 14 and 28 days of exposure to 6 mL/kg (Fig. S2).

### Resting metabolic rate

To measure RMR, we removed birds from their cages 30 minutes after the aviary lights turned off at 2100 h and placed birds in 1 L respirometry jars supplied with air at a flow rate of 400 ml/minute. Respirometry jars were placed inside an environmental chamber and temperature was maintained at 28 C. Air was scrubbed of CO_2_ and water vapor using reconstituted Drierite columns prior to entering the respirometry chambers. One hour after being transferred to the respirometry jars, we measured CO_2_ production and O_2_ consumption using a Field Metabolic System gas analyzer (Sable Systems). RMR data was collected for no more than 4 birds at a time. Each respirometry jar was sampled for 3 minutes every 15 minutes using a Flow Bar 8 and Multiplexer (Sable Systems). Baseline O_2_ and CO_2_ measurements (i.e., flow through an empty respirometry jar) was also measured for 3 minutes every 15 minutes to correct for drift. Birds were returned to their cages 2 hours before the aviary lights came on at 0430 h. To calculate RMR, we first calculated VO_2_ and VCO_2_ using equations described by Bartholomew et al. (1981). We then calculated mean VO_2_ for each sampling interval, and RMR was defined as the 3-minute sampling period with the lowest mean VO_2_.

### Maximum metabolic rate

To measure MMR, we constructed an air-tight hop-flutter wheel (diam.= 35 cm; height=14 cm) using a polypropylene cylinder with plexiglass panels affixed to both sides (sensu Chappell et al. 2006^8^). The hop-flutter wheel was powered by a 12 V DC motor (MoterMaker Inc.), and a potentiometer facilitated manual control of the rotational speed of the wheel. Air was scrubbed of CO_2_ and water vapor using Drierite and pumped into the hop-flutter wheel at a flow rate of 4200 ml/min using a Flow Bar 8. Excurrent air was subsampled (200 ml/min) and O_2_ consumption and CO_2_ production were measured on a Field Metabolic System. At the start of each trial, we collected 5 min of baseline gas concentrations, then a single bird was placed in the hop-flutter wheel for 5 min without rotation. The hop-flutter wheel was covered during this initial reading. We then uncovered the wheel and started to slowly increase rotational speed of the hop-flutter chamber until finches continuously hopped or attempted to hover inside the wheel. Rotational speed was adjusted for each bird so that the birds was constantly hopping/fluttering, and varied between 35 and 45 rpm. %O_2_ in excurrent air was measured in real-time, and birds were exercised until the O_2_ consumption profile plateaued and further increasing the rotation of the hop-flutter chamber did not increase O_2_ consumption, at which point the hop-flutter wheel was turned off. All birds were visibly panting after the hop-flutter wheel was turned off. Birds typically achieved MMR within 150 s from when rotation was initiated. After birds were removed from the chamber, we collected 3 additional min of baseline gas concentrations to correct for any drift during the trial. All MMR trials were conducted between 1100 h and 1200 h.

### Flight chamber construction & vertical flight speed

To measure vertical speed, we constructed a vertical take-off chamber similar to Kullberg et al. 2002^9^. The chamber consisted of a PVC frame and white poster-board that formed a semi-circle around the PCV frame. We affixed a clear plexiglass panel to the open side of the poster-board semi-circle so that birds were visible for the entirety of their flight. At the top of the chamber was a cardboard platform covering the vertical flight chamber with 18×18 cm square hole, and a cage (30×40×40 cm) with wooden perches was placed on the cardboard platform over the square hole. During a flight trial, birds would fly through the square hole at the top of the flight chamber and land on a wooden perch. A second cardboard platform was placed at the bottom of the flight chamber, upon which we secured a wooden perch. The final dimensions of the vertical flight chambers were height = 230 cm and radius = 22 cm.

Prior to the experiment, we collected morphometric data for each bird (Table S2). Each vertical flight attempt was video-recorded using a Sony camcorder that captured video at 28-30 frames per second (fps) placed 4 m from the flight chamber. To eliminate the potential effect of birds slowing down as they approached the top platform, we excluded the top 46 cm of the flight chamber from the analysis. Similarly, we did not include the bottom 46 cm to account for variation in take-off performance. Therefore, the functional distance analyzed was 138 cm. Videos were analyzed using ImageJ, whereby we converted videos to individual frames and digitally measured the distance from where the bird first crossed the bottom threshold to just before the bird crossed the top threshold. All birds were within 1 body length of both the top and bottom thresholds. Speed was calculated by dividing the measured linear distance traveled between the bottom and top threshold by the time it to took a bird to travel this distance. We only calculated speed for successful trials, when birds flew into the cage above the flight chamber. For each bird, we calculated the mean flight speed across four successful attempts. A single observer (JBK) who was blind to treatment analyzed all flight speed videos.

---

**Table S1:** Summary of hypotheses and predictions tested in the present study.

| HYPOTHESES | MEASURED RESPONSE | PREDICTED OUTCOME |
| --- | --- | --- |
| (1) Crude oil ingestion will cause oxidative damage to RBCs and a shift in hematological indices | Oxidative damage (HDP fluorescence) | ↑ |
|  | PCV & Hb conc. | ↓ |
|  | Reticulocytes | ↑ |
| (2) Shifts in hematological indices will interfere with blood-oxygen transportation and cause a reduction in aerobic scope for activity | RMR | ↑ |
|  | MMR | ↓ |
|  | Aerobic scope | ↓ |
| (3) A reduction in aerobic scope for activity will impair flying performance | Short distance flight speed | ↓ |

**Table S2:** Morphometric measurements (mm) of experimental birds included in flight trials.

| | Control<br>(mean ± std. dev.) | 3 mL/kg<br>(mean ± std. dev.) | 6 mL/kg<br>(mean ± std. dev.) | *F* | *p* |
| --- | --- | --- | --- | --- | --- |
| Tarsus | 14.41 ± 0.75 | 13.80 ± 0.59 | 13.64 ± 0.86 | 2.25 | 0.14 |
| Wing Chord | 64.25 ± 7.48 | 65.14 ± 3.39 | 63.00 ± (2.76) | 0.27 | 0.77 |
| Wing Length | 61 ± 5.48 | 60.43 ± 1.72 | 59.83 ± 1.60 | 0.18 | 0.84 |
| Body Width | 17.84 ± 0.02 | 18.76 ± 1.34 | 19.39 ± 2.11 | 0.95 | 0.41 |

**Table S3:** PAH concentrations in artificially weathered (85% original volume) crude oil.

| Analyte | Abb. | mg/kg | TPAH50 |
| --- | --- | --- | --- |
| Naphthalene | N0 | 480 | X |
| C1 - Naphthalenes | N1 | 1600 | X |
| C2 - Naphthalenes | N2 | 2200 | X |
| C3 - Naphthalenes | N3 | 1600 | X |
| C4 - Naphthalenes | N4 | 800 | X |
| Biphenyl | B | 250 | X |
| Acenaphthylene | AY | 18 | X |
| Acenaphthene | AE | 17 | X |
| Fluorene | F0 | 160 | X |
| C1 - Fluorenes | F1 | 340 | X |
| C2 - Fluorenes | F2 | 460 | X |
| C3 - Fluorenes | F3 | 390 | X |
| Anthracene | A0 | 16 | X |
| Phenanthrene | P0 | 280 | X |
| C1 - Phenanthrenes/Anthracenes | PA1 | 690 | X |
| C2 - Phenanthrenes/Anthracenes | PA2 | 810 | X |
| C3 - Phenanthrenes/Anthracenes | PA3 | 550 | X |
| C4 - Phenanthrenes/Anthracenes | PA4 | 310 | X |
| Retene | T | 20 | X |
| Dibenzothiophene | DBT0 | 38 | X |
| C1 - Dibenzothiophenes | DBT1 | 120 | X |
| C2 - Dibenzothiophenes | DBT2 | 200 | X |
| C3 - Dibenzothiophenes | DBT3 | 150 | X |
| C4 - Dibenzothiophenes | DBT4 | 78 | X |
| Fluoranthene | FL0 | 5.2 | X |
| Pyrene | PY0 | 14 | X |
| C1 - Fluoranthenes/Pyrenes | FP1 | 45 | X |
| C2 - Fluoranthenes/Pyrenes | FP2 | 60 | X |
| C3 - Fluoranthenes/Pyrenes | FP3 | 48 | X |
| C4 - Fluoranthenes/Pyrenes | FP4 | 36 | X |
| Naphthobenzothiophene | NBT0 | 10 | X |
| C1 - Naphthobenzothiophenes | NBT1 | 95 | X |
| C2 - Naphthobenzothiophenes | NBT2 | 160 | X |
| C3 - Naphthobenzothiophenes | NBT3 | 180 | X |
| C4 - Naphthobenzothiophenes | NBT4 | 130 | X |
| Benz(a)anthracene | BA0 | 11 | X |
| Chrysene | C0 | 47 | X |
| C1 - Chrysenes | BC1 | 120 | X |
| C2 - Chrysenes | BC2 | 160 | X |
| C3 - Chrysenes | BC3 | 120 | X |
| C4 - Chrysenes | BC4 | 95 | X |
| Benzo(a)fluoranthene | BAF | ND | X |
| Benzo(j+k)perylene | BJKF | ND | X |
| Benzo(b)fluoranthene | BAF | 5.9 | X |
| Benzo(e)pyrene | BEP | 8.5 | X |
| Benzo(a)pyrene | BAP | ND | X |
| Perylene | PER | 14 | X |
| Indeno(1,2,3-cd)pyrene | IND | ND | X |
| Dibenzo(a,h)anthracene | DA | ND | X |
| Benzo(g,h,i)perylene | GHI | ND | X |
| 1-Methylnaphthalene | NM1 | 920 |  |
| 1-Methylphenanthrene | PM1 | 210 |  |
| 2-Methylantracene | AM1 | 11 |  |
| 2-Methyldibenzothiophene | DBTM2 | 34 |  |
| 2-Methylnaphthalene | NM2 | 1700 |  |
| 2-Methylphenanthrene | PM1 | 170 |  |
| 2,3,5-Trimethylnaphthalene | N3M | 430 |  |
| 2,6-Dimethylnaphthalene | N2M | 1100 |  |
| 3-Methylphenanthrene | PM3 | 160 |  |
| 4-Methyldibenzothiophene | DBTM3 | 65 |  |
| 9-Methylphenanthrene | PM9 | 200 |  |
| Benzo(b)fluorene | BF | 15 |  |
| Benzo(b)thiophene | BBT | 8.3 |  |
| C1 - Benzothiophenes | BT1 | 43 |  |
| C1 - Decalins | D1 | 710 |  |
| C2 - Benzothiophenes | BT2 | 39 |  |
| C2 - Decalins | D2 | 1200 |  |
| C3 - Benzothiophenes | BT3 | 29 |  |
| C3 - Decalins | D3 | 1200 |  |
| C30-Hopane | H30 | 93 |  |
| C4 - Benzothiophenes | BT4 | 32 |  |
| C4 - Decalins | D4 | 1200 |  |
| Carbazole | CB | 4.1 |  |
| Decahydronaphthalene, Total | DHPT | 320 |  |
| Dibenzofuran | DF | 29 |  |

**Fig. S1:**
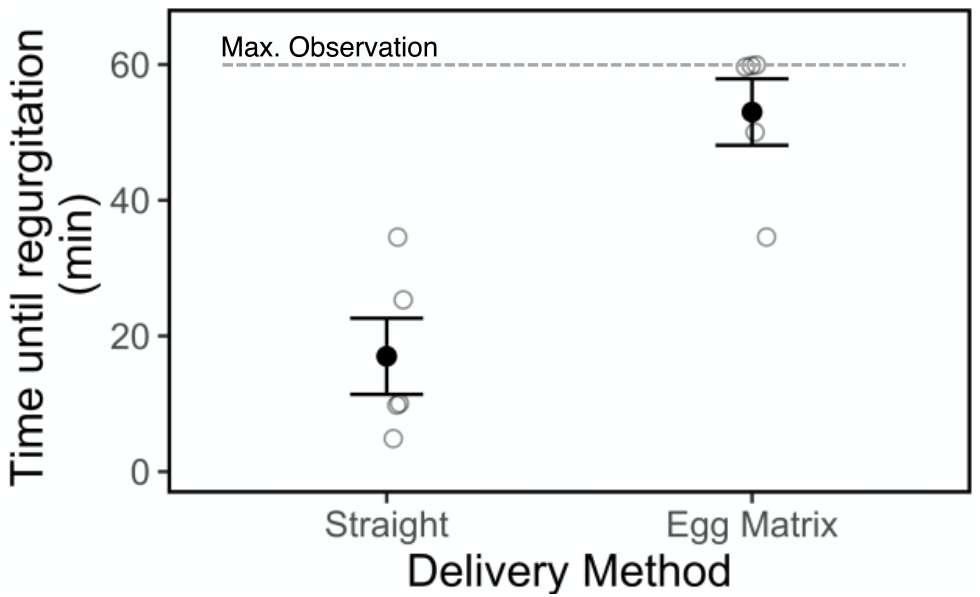
Regurgitation in zebra finches dosed with a single dose of 3 mL/kg bird mass straight or within an egg matrix. Birds were monitored individually for up to 60 min.

**Fig. S2:**
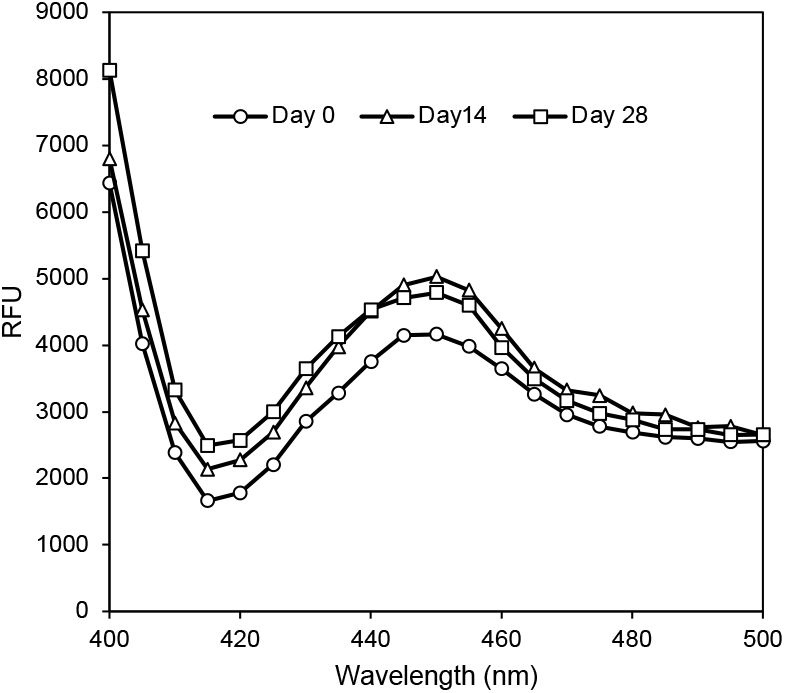
Representative heme degradation product emission spectra of whole blood from the same zebra finch prior to exposure (day 0) and after 14 and 28 days of exposure to 6 mL/kg.

**Fig. S3:**
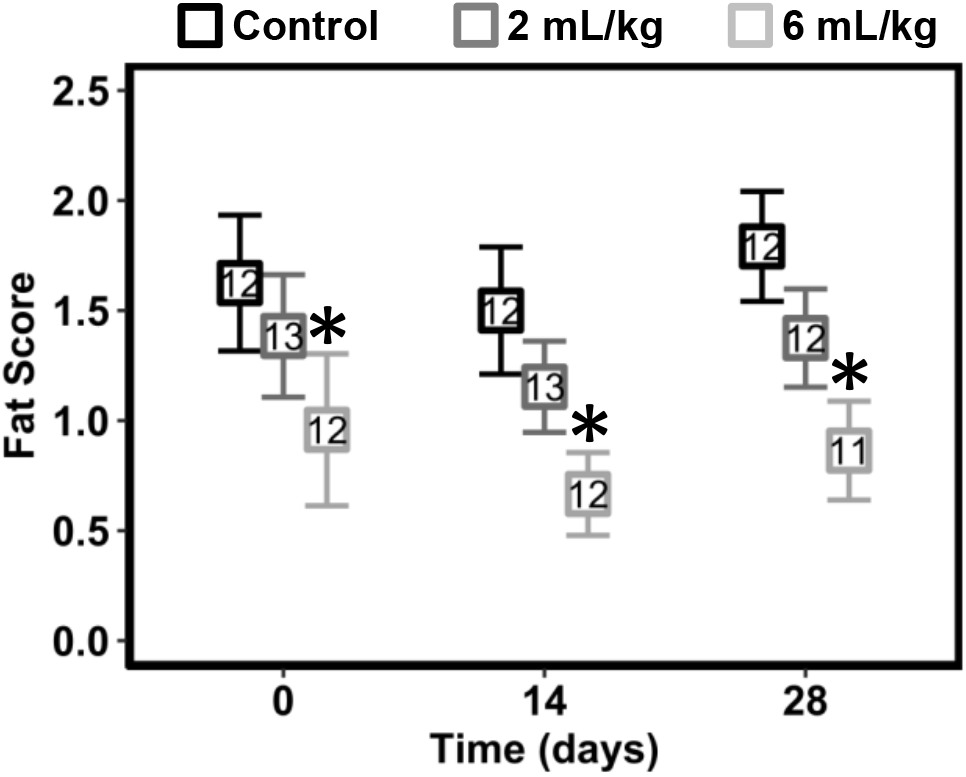
Fat scores of zebra finches prior to dosing (day 0) and after 14 and 28 days of oral dosing with peanut oil (controls) or weathered crude oil (squares and whiskers represent mean±std. error, numbers within squares represent sample sizes, asterisks denote significant difference [p<0.05] compared to the control).

